# Tomato CYP94C1 terminates jasmonate signaling during fruit ripening by inactivating bioactive jasmonoyl-L-isoleucine

**DOI:** 10.1101/2023.05.17.541065

**Authors:** Tianxia Yang, Lei Deng, Chuanlong Sun, Muhammad Ali, Fangming Wu, Huawei Zhai, Qian Xu, Peiyong Xin, Jinfang Chu, Tingting Huang, Chang-Bao Li, Chuanyou Li

**Affiliations:** State Key Laboratory of Plant Genomics, National Center for Plant Gene Research (Beijing), Institute of Genetics and Developmental Biology, Innovation Academy for Seed Design, Chinese Academy of Sciences, Beijing 100101, China; CAS Center for Excellence in Biotic Interactions, University of Chinese Academy of Sciences, Beijing 100049, China; State Key Laboratory of Crop Biology, College of Horticulture Science and Engineering, College of Life Sciences, Shandong Agricultural University, Tai’an 271018, China; Institute of Vegetable, Qingdao Academy of Agricultural Sciences, Qingdao 266100, China; Key Laboratory of Biology and Genetic Improvement of Horticultural Crops (North China), Ministry of Agriculture, Beijing Vegetable Research Center, Beijing Academy of Agriculture and Forestry Sciences, Beijing 100097, China

**Keywords:** tomato, defense, ripening, jasmonate, ethylene, *CYP94*, transcription regulation

## Abstract

Ripe fruits are more susceptible to necrotrophic pathogens than unripe fruits. Although this phenomenon is widespread across different fruit species and results in substantial economic losses, the underlying mechanism is still poorly understood. Previous studies revealed that ethylene (ET) is a key signal controlling climacteric fruit ripening and that jasmonate (JA) regulates plant resistance to necrotrophs. We investigated the function of tomato cytochrome P450 94 (*CYP94*) family genes in JA signaling and report here that ET-mediated ripening suppresses JA-mediated defense by promoting the deactivation of bioactive JA-Ile. ETHYLENE-INSENSITIVE 3 (EIN3)/EIN3-LIKE (EIL) transcription factors directly activated *CYP94C1* to convert JA-Ile to its inactive form 12-COOH-JA-Ile, thereby terminating JA signaling during fruit ripening. Mutation of *CYP94C1* led to increased resistance of ripe fruits to the necrotrophic pathogen *Botrytis cinerea* without affecting the ripening process. Additionally, the master transcription factor MYC2 directly activated two other CYP94 members *CYP94B1* and *CYP94B2* to convert JA-Ile to its less active form 12-OH-JA-Ile, thereby attenuating JA signaling in wounded leaves. Simultaneous mutation of *CYP94B1* and *CYP94B2* increased the resistance of leaves to *B. cinerea*. Thus, differences in the expression and enzymatic activities of *CYP94* family gene members precisely control JA-mediated defense responses in tomato.

## INTRODUCTION

Fruit rot caused by necrotrophic pathogens results in substantial reductions in fruit yield and revenues worldwide (Petrasch et al., 2019b; Prusky et al., 2013; Silva et al., 2021). A widespread phenomenon in fleshy fruit species is the susceptibility of ripe fruits to necrotrophs (Alkan and Fortes, 2015; Balsells-Llaurado et al., 2020; Blanco-Ulate et al., 2016; Cantu et al., 2009; Forlani et al., 2019; Haile et al., 2019; Li et al., 2022; Petrasch et al., 2019a; Petrasch et al., 2019b; Silva et al., 2023; Silva et al., 2021), which facilitates seed dispersal (Forlani et al., 2019), but causes severe postharvest losses in production. As most of the nutritional and sensory qualities of fruits are elaborated at the ripening stage (Klee and Giovannoni, 2011; Liu et al., 2015), balancing fruit ripening and pathogen resistance to maintain fruit quality has proven to be challenging (Adaskaveg and Blanco-Ulate, 2023; Li et al., 2022). A deeper understanding of the mechanisms underlying the increased susceptibility of fruits to necrotrophs during ripening could lead to new strategies for producing necrotrophy-resistant fruits without compromising ripening-related quality.

Fruit ripening is a highly coordinated developmental process that involves a wide range of changes in color, texture, flavor, aroma, and other quality attributes (Chen et al., 2020; Klee and Giovannoni, 2011; Liu et al., 2015). Many of these changes have been suggested or shown to contribute to susceptibility to necrotrophs (Blanco-Ulate et al., 2016). First, the disassembly of the cell wall during fruit softening renders it ineffective as a physical barrier to pathogens (Cantu et al., 2008; Prusky et al., 2013; Silva et al., 2021). Second, alterations in metabolite contents (such as sugars, amino acids, and organic acids) and apoplastic pH may provide a favorable environment for pathogen invasion and/or colonization (Blanco-Ulate et al., 2016; Li et al., 2022; Manteau et al., 2003; Petrasch et al., 2019a). Third, constitutive and inducible defenses against necrotrophs decline during fruit ripening (Beno-Moualem and Prusky, 2000; Haile et al., 2019; Prusky et al., 2013; You and van Kan, 2021), although the underlying molecular details remain largely unknown.

Ethylene (ET) is a key signal that controls most aspects of ripening in climacteric fruits (Chen et al., 2020; Fenn and Giovannoni, 2021; Klee and Giovannoni, 2011). ETHYLENE-INSENSITIVE 3 (EIN3)/EIN3-LIKE (EIL) transcription factors are at the core of the ET signaling pathway (Chao et al., 1997; Merchante et al., 2013; Tieman et al., 2001; Yokotani et al., 2009). In tomato (*Solanum lycopersicum*), the most studied climacteric fruit, EIL regulates the expression of ripening-related regulatory genes as well as structural genes through promoter binding (Deng et al., 2023; Huang et al., 2022; Lu et al., 2018). Mutation of its downstream regulators (e.g. *NON-RIPENING* [*NOR*]) and cell wall disassembly genes (e.g. *PECTATE LYASE* [*PL*]) leads to increased resistance to the necrotrophic pathogen *Botrytis cinerea* in ripe fruits (Cantu et al., 2009; Cantu et al., 2008; Silva et al., 2021), suggesting a negative role for ET in regulating pathogen resistance during ripening.

Jasmonate (JA) is a major defense hormone that promotes plant resistance to mechanical wounding, chewing insects, and necrotrophic pathogens (Glazebrook, 2005; Howe and Jander, 2008). The basic helix-loop-helix (bHLH) transcription factor MYC2 acts as the master regulator of diverse aspects of JA responses (Boter et al., 2004; Du et al., 2017; Kazan and Manners, 2013; Lorenzo et al., 2004; Zhai et al., 2020). Jasmonoyl-l-isoleucine (JA-Ile) is the most bioactive form of JA (Fonseca et al., 2009; Katsir et al., 2008; Sheard et al., 2010; Thines et al., 2007; Yan et al., 2009). In the absence of JA-Ile, JASMONATE-ZIM DOMAIN (JAZ) proteins interact with and repress the transcriptional activity of MYC2 (Chini et al., 2007; Thines et al., 2007). In response to stress or developmental cues, elevated JA-Ile levels promote the degradation of JAZ repressors, thereby activating MYC2-directed transcription of JA-responsive genes (Chini et al., 2007; Fonseca et al., 2009; Sheard et al., 2010; Thines et al., 2007).

JA-Ile is deactivated by the ω-oxidation pathway, in which it is converted to its less active form 12-hydroxy-JA-Ile (12-OH-JA-Ile) and then further oxidized to its inactive form 12-dicarboxy-JA-Ile (12-COOH-JA-Ile) (Aubert et al., 2015; Heitz et al., 2012; Kitaoka et al., 2011; Koo et al., 2011; Koo and Howe, 2012; Koo et al., 2014). In *Arabidopsis*, this pathway is mediated by three cytochrome P450 oxidases, CYP94B1, CYP94B3, and CYP94C1. While CYP94B1 and CYP94B3 preferentially hydroxylate JA-Ile to 12-OH-JA-Ile, CYP94C1 further carboxylates 12-OH-JA-Ile to 12-COOH-JA-Ile (Aubert et al., 2015; Heitz et al., 2012; Kitaoka et al., 2011; Koo et al., 2011; Koo and Howe, 2012; Koo et al., 2014). Interestingly, all three *CYP94* genes are induced by wounding (Heitz et al., 2012; Koo et al., 2011) and by *B. cinerea* infection (Aubert et al., 2015). Thus, they can exert negative feedback control of JA-Ile levels, providing an efficient route for the attenuation or termination of JA signaling.

Although the role of JA in regulating resistance to necrotrophic pathogens has been intensively studied in leaves (Du et al., 2017; Thomma et al., 1998; Wasternack and Song, 2017; Yan et al., 2013; Zhu et al., 2011), less is known about its roles in fruits, especially during the ripening process. In addition, although JA and ET synergistically promote resistance in leaves, they likely have opposing roles in ripe fruits (Blanco-Ulate et al., 2016; Lorenzo et al., 2004; Lorenzo et al., 2003; Silva et al., 2021; Zhu et al., 2011), suggesting a complex interplay between these two hormones. The relationship between ET and JA signaling during fruit ripening and its impact on pathogen resistance remain to be elucidated.

Here, we investigated the function of tomato *CYP94* family genes in JA signaling and found that ET-mediated ripening suppresses JA-mediated defense by deactivating JA-Ile. EIL directly activated the expression of *CYP94C1* to convert JA-Ile to its inactive form 12-COOH-JA-Ile, terminating JA signaling during fruit ripening. Knockout of *CYP94C1* specifically increased the resistance of ripe fruits to *B. cinerea* without affecting the ripening process. Additionally, MYC2 directly activated the expression of *CYP94B1* and *CYP94B2* to convert JA-Ile to its less active form 12-OH-JA-Ile, thereby attenuating JA signaling in wounded leaves. Mutation of *CYP94B1* and *CYP94B2* led to increased resistance of leaves to *B. cinerea*. Therefore, differences in the expression and enzymatic activities of *CYP94* family genes precisely control JA-mediated defense responses in tomato.

## RESULTS

### JA-Ile level and JA-mediated defense responses decline during fruit ripening

Given the crucial role of ET and JA in regulating ripening and resistance to necrotrophs, respectively, we monitored the levels of ET and JA-Ile in the ripening process of wild-type (WT) fruits. As expected, ET production was negligible before the onset of ripening (i.e., at the mature green [MG] stages) but was highly induced upon initiation of ripening (i.e., at the breaker [Br] stage and 4 days after the Br [Br+4] stage) (Figure 1A). By contrast, JA-Ile content was high before the onset of ripening but was markedly reduced upon ripening initiation (Figure 1A).

**Figure 1.**
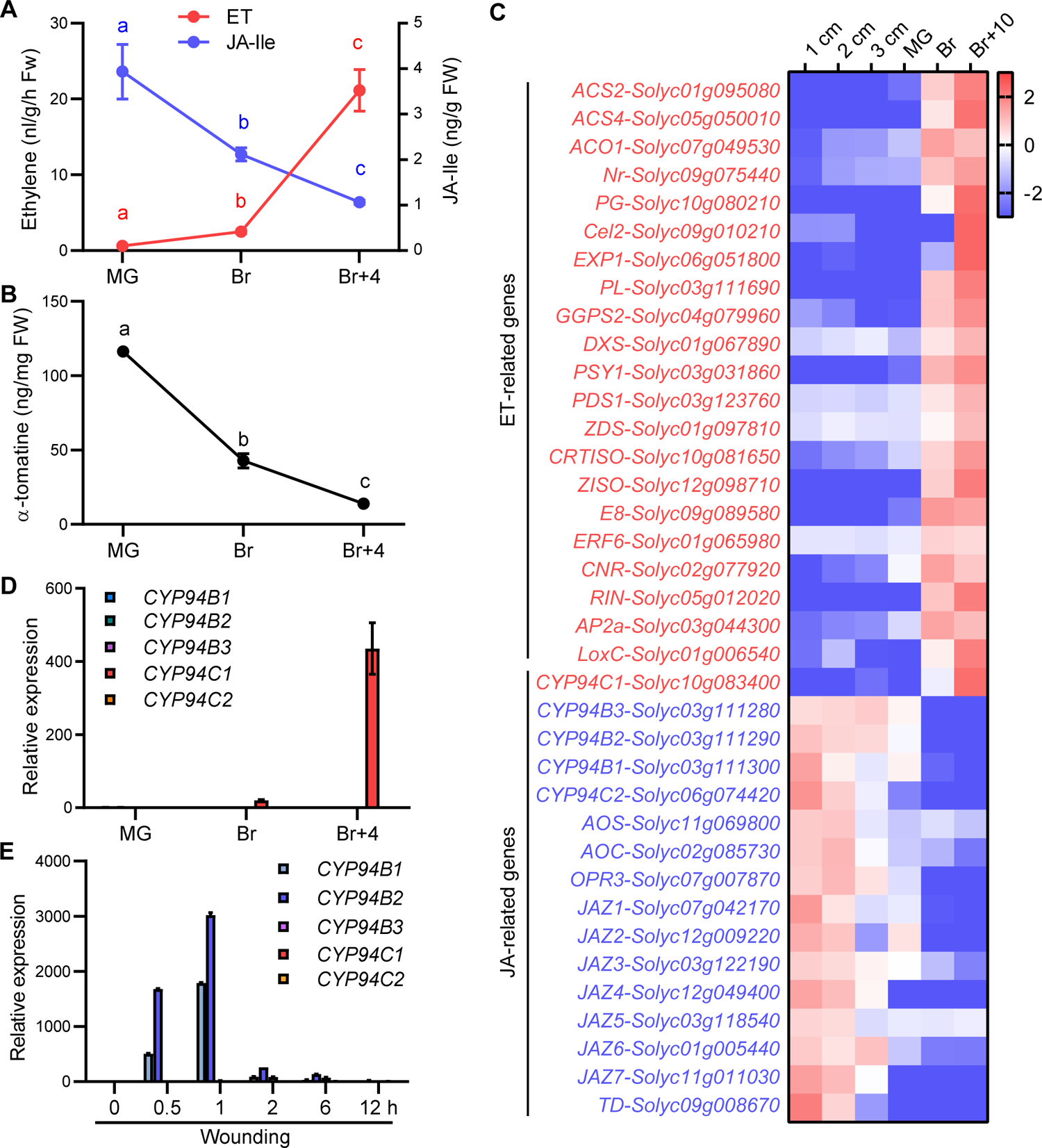
JA-Ile level and JA-mediated defense responses are decreased during fruit ripening. **(A)** ET and JA-Ile levels during fruit ripening. Data are mean ± SD, *n* = 3 repeats. **(B)** α-tomatine level during fruit ripening. Data are mean ± SD, *n* = 3 repeats. **(C)** Expression profiles of well-characterized ET- and JA-related genes. Expression data were obtained from the Tomato Expression Database (http://ted.bti.cornell.edu/). The scale bar represents log_2_(fold change [FC]) of the average expression level of each gene in three biological replicates. MG, mature green stage; Br, breaker stage; Br+10, 10 days after the Br stage. **(D)** RT-qPCR results showing the expression levels of five *CYP94* genes in the tomato fruit at different developmental stages. Data are mean ± SD, *n* = 3 repeats. **(E)** RT-qPCR results showing wound-induced expression of five *CYP94* genes in tomato leaves. Data are mean ± SD, *n* = 3 repeats. For **(A and B)**, fruits were harvested at the MG (i.e., 39 days post anthesis [DPA]), Br (i.e., 42 DPA), and Br+4 (i.e., 46 DPA) stages. The bars with different letters are significantly different from each other (*P* < 0.05; Student’s *t* test).

To investigate how JA-mediated defense changes during fruit ripening, we first examined the level of α-tomatine, a defensive secondary metabolite, the biosynthesis of which is positively regulated by the JA pathway (Cardenas et al., 2016; Panda et al., 2022). Unsurprisingly, consistent with the decline in the JA level, α-tomatine decreased dramatically during fruit ripening (Figure 1B). We then examined the expression profiles of JA- and ET-related genes during ripening using the public Tomato Expression Database (TED; http://ted.bti.cornell.edu/) (The Tomato Genome Consortium, 2012). As expected, many well-characterized ET-related genes (Deng et al., 2023) were induced by ripening (Figure 1C), including several ET biosynthetic and signaling genes, several ripening-related regulators, and various ripening-related structural genes (Figure 1C). By contrast, many well-characterized JA-related genes (Du et al., 2017) were repressed by ripening, including several JA biosynthetic genes, such as *ALLENE OXIDE SYNTHASE* (*AOS*), *ALLENE OXIDE CYCLASE* (*AOC*), and *12-OXOPHYTODIENOATE REDUCTASE3* (*OPR3*), several *JAZ* genes, and the defense gene *THREONINE DEAMINASE* (*TD*) (Figure 1C). These results indicate that JA-mediated defense responses also decrease during ripening.

We also examined JA-Ile and α-tomatine levels in RNAi-mediated *EIL* knockdown (*EIL-RNAi*) fruits, which exhibit severe defects in autocatalytic ET biosynthesis and fruit ripening (Deng et al., 2023). Ripening-induced decreases in JA-Ile and α-tomatine levels were significantly slower to increase in *EIL-RNAi* fruits than in WT fruits (Supplemental Figures 1A and 1B), suggesting that the decline of JA-Ile and JA-mediated defense responses is an integral part of fruit ripening.

To understand why the level of JA-Ile declines during ripening, we examined the expression pattern of tomato *CYP94* gene family members. Homology analysis indicated that the tomato genome harbors five *CYP94* genes: *CYP94B1*, *CYP94B2*, *CYP94B3*, *CYP94C1*, and *CYP94C2* (Supplemental Figures 2A and 2B). Among them, *CYP94B1*, *CYP94B2*, and *CYP94B3* cluster together on chromosome 3 (Supplemental Figure 2C). Notably, transcriptome data showed that while the expression of *CYP94B1*, *CYP94B2*, *CYP94B3*, and *CYP94C2* was repressed by ripening, the expression of *CYP94C1* was induced by ripening (Figure 1C). To validate these results, we performed reverse transcription quantitative PCR (RT-qPCR) assays using WT fruits harvested at different developmental stages. *CYP94C1* expression was low before the onset of ripening (i.e., at the MG stage) but was highly induced after the onset of ripening (i.e., at the Br and Br+4 stages) (Figure 1D). By contrast, *CYP94B1*, *CYP94B2*, *CYP94B3*, and *CYP94C2* showed negligible expression in both unripe and ripe fruits (Figure 1D). We also examined the temporal expression patterns of these genes in response to mechanical wounding. RT-qPCR assays revealed that the expression of *CYP94B1* and *CYP94B2*, but not that of other *CYP94* genes, was highly induced by wounding (Figure 1E). Collectively, these results demonstrated that *CYP94C1* is a ripening-induced gene, while *CYP94B1* and *CYP94B2* are wound-induced genes. Taken together, the above findings led us to hypothesize that CYP94C1 converts JA-Ile to its inactive or less active form in ripe fruits, thereby reducing the level of JA-Ile and JA-mediated defense responses during ripening.

**Figure 2.**
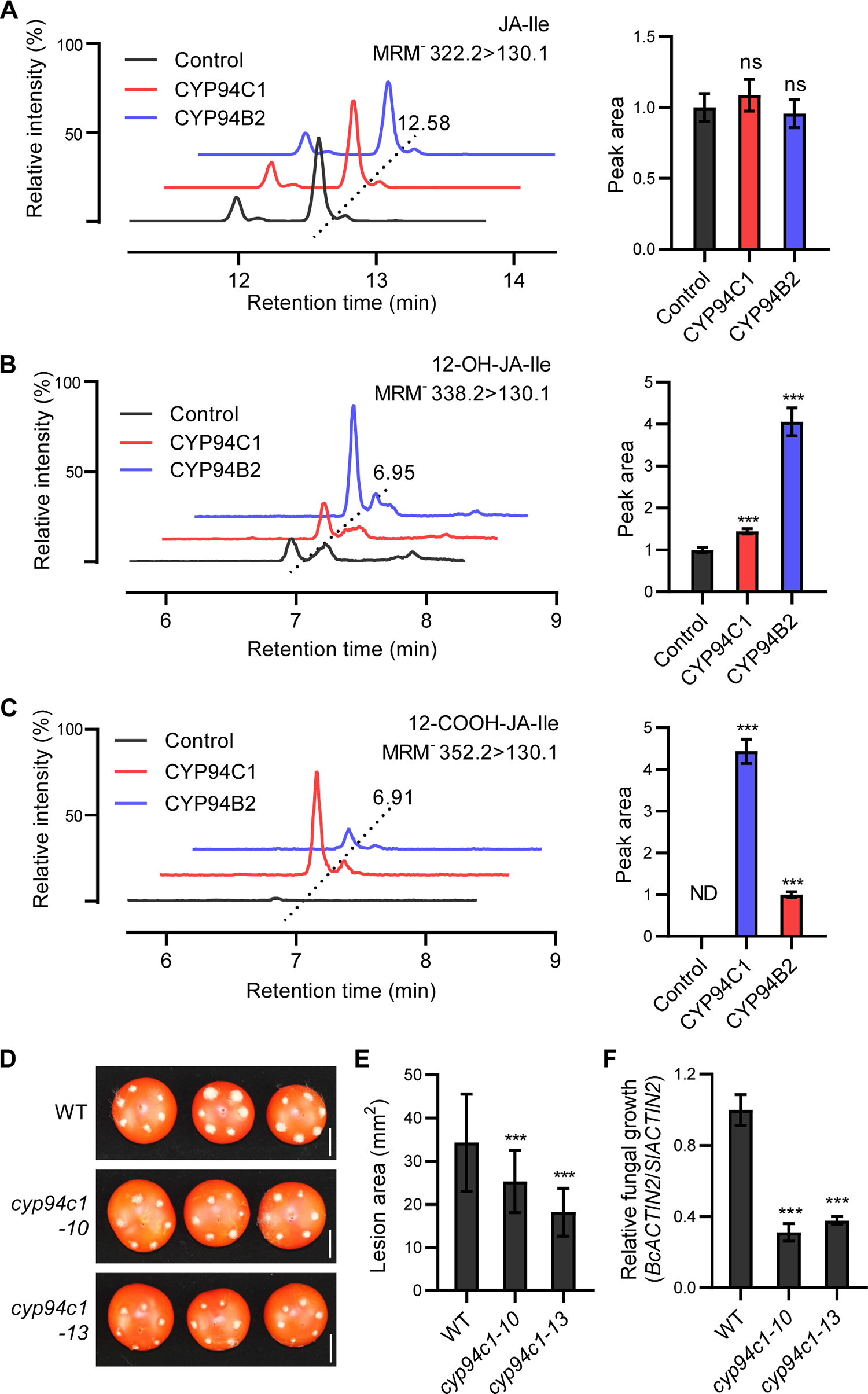
CYP94C1 negatively regulates ripe fruit resistance to *B. cinerea* by converting JA-Ile to the inactive 12-COOH-JA-Ile. **(A–C)** LC chromatogram (left panel) of residual JA-Ile **(A)**, 12-OH-JA-Ile **(B)**, and 12-COOH-JA-Ile **(C)** produced in incubations of liquid culture of yeast cells expressing *CYP94B2* or *CYP94C1*. Reaction products were identified by LC-MS/MS based on their retention times and detection in multiple reaction monitoring (*MRM*, *m/z* 322.2 > 130.1 for JA-Ile, *m/z* 338.2 > 130.1 for 12-OH-JA-Ile, and *m/z* 352.2 > 130.1 for 12-COOH-JA-Ile). The right panel shows the relative peak area of indicated metabolites. Data are mean ± SD, *n* = 3 repeats. Asterisks indicate significant*** differences between control and CYP94B2 or CYP94C1 ( *P* < 0.001; Student’s *t* test). ns, not significant; ND, not detected. **(D)** Representative images of *B. cinerea*-inoculated WT and *cyp94c1* fruits. Fruits harvested at the Br+4 stage were inoculated with *B. cinerea*. The images were taken 3 days post inoculation. Bars = 2 cm. **(E)** Lesion area on WT and *cyp94c1* fruits. Data are mean ± SD, *n* = 27 repeats. **(F)** Quantification of the fungal growth on WT and *cyp94c1* fruits. Data are mean ± SD, *n* = 3 repeats. For **(E and F)**, asterisks indicate significant differences between WT and *cyp94c1*\*\*\* mutants ( *P*<0.001; Student’s *t* test).

### CYP94C1 terminates JA signaling in ripe fruits by converting JA-Ile to inactive 12-COOH-JA-Ile

To test the above hypothesis, we first examined the enzymatic activity of CYP94C1 by using a yeast expression system specifically developed for P450 (Nomura et al., 2013; Pompon et al., 1996). Liquid chromatography-tandem mass spectrometry (LC-MS/MS) analysis detected high levels of 12-COOH-JA-Ile and relatively low levels of 12-OH-JA-Ile after incubation of JA-Ile with the liquid culture of yeast cells expressing *CYP94C1* (Figures 2A–2C), suggesting that CYP94C1 catalyzes two successive ω-oxidation steps of JA-Ile and tends to convert JA-Ile to its inactive form 12-COOH-JA-Ile, as described previously for its *Arabidopsis* ortholog (Heitz et al., 2012).

We then generated null alleles of *CYP94C1* by clustered regularly interspaced short palindromic repeats/CRISPR-associated 9 (CRISPR/Cas9)-mediated gene editing (Cong et al., 2013; Mali et al., 2013). Two resulting mutants (c*yp94c1-10* and *cyp94c1-13*), each carrying a mutation causing a frameshift and premature termination of translation (Supplemental Figures 3A–3C), were selected for further analysis. We first examined the properties of their ripe fruits after *B. cinerea* infection. As shown in Figures 2D and 2E, the size of the necrotic lesion on *cyp94c1* fruits caused by *B. cinerea* was much larger than that on WT fruits. Consistently, fungal growth, as assessed by qPCR amplification of the *B. cinearea ACTIN2* gene (*BcACTIN2*) relative to that of the tomato *ACTIN2* gene (*SlACTIN2*), was significantly higher on *cyp94c1* fruits than on WT fruits (Figure 2F). These results indicate that *cyp94c1* fruits are more resistant to *B. cinerea* than WT fruits. Notably, mutant fruits initiated ripening at a similar time to WT fruit, suggesting that the depletion of *CYP94C1* does not impair the ripening process (Supplemental Figure 3D). Additionally, we examined the properties of their leaves after *B. cinerea* infection. The lesion size and fungal growth on *cyp94c1* leaves were comparable with those on WT leaves, indicating that knockout of *CYP94C1* does not affect leaf resistance to *B. cinerea* (Supplemental Figures 3E–3G). Collectively, these results suggest that CYP94C1 negatively regulates JA-mediated defense responses in ripe fruits.

**Figure 3.**
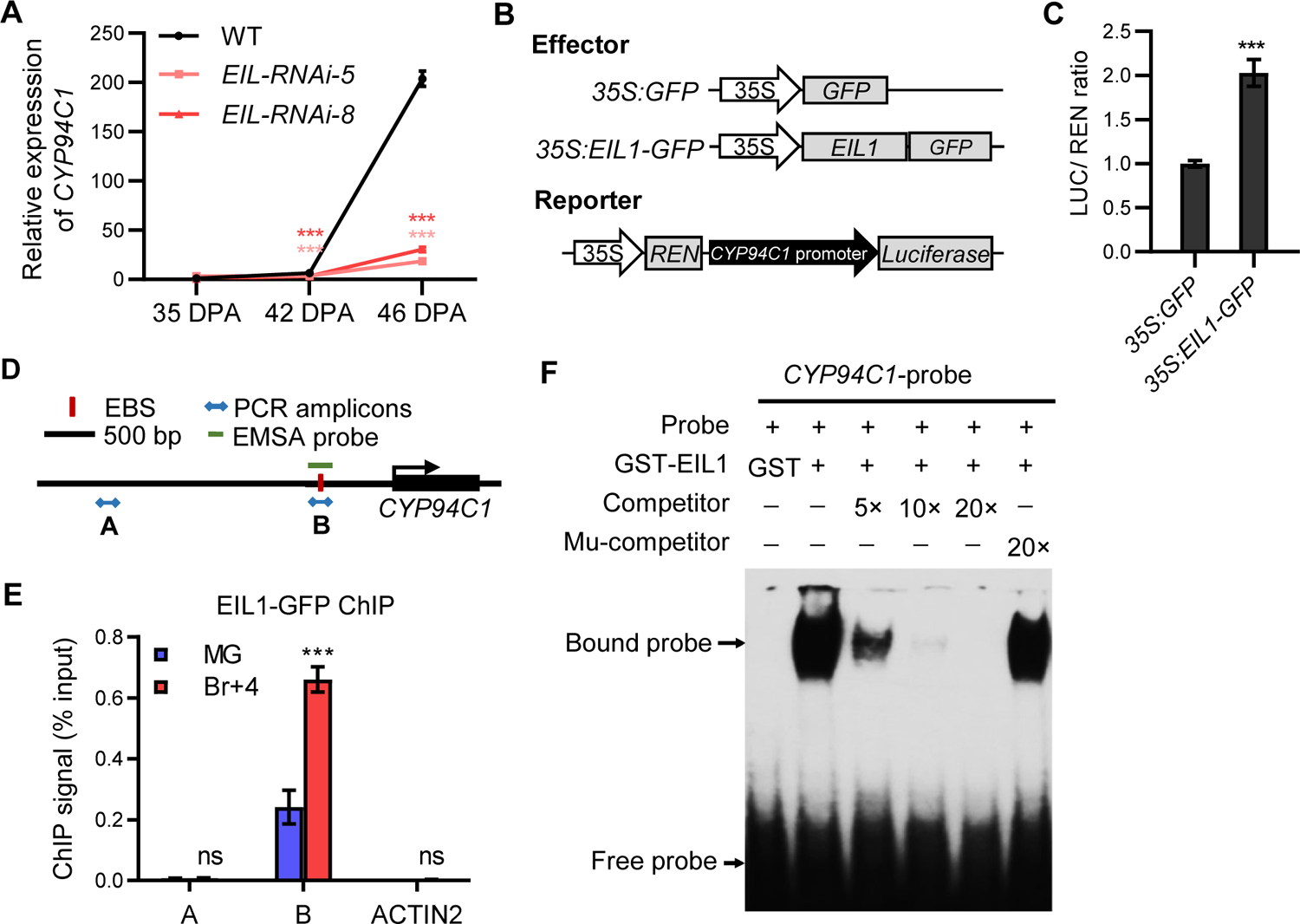
EIL regulates *CYP94C1* expression by binding to its promoter. **(A)** RT-qPCR results showing the expression of *CYP94C1* in WT and *EIL-RNAi* fruits. Fruits were harvested at 35, 42, and 46 DPA, which corresponds to the MG stage, Br stage, and Br+4 stage in the WT, respectively. Data are mean ± SD, *n* = 3 repeats. Asterisks indicate significant differences between WT and *EIL-RNAi* fruits (\*\*\**P*<0.001; Student’s *t* test). **(B)** Schematic diagram showing the constructs used in the transient expression assays of **(C)**. Arrows, promoter regions; shaded boxes, coding regions. **(C)** Transient expression assays of *N. benthamiana* leaves showing activation of the *CYP94C1* promoter by EIL1. The *ProCYP94C1:LUC* reporter was co-transfected with the indicated effector constructs. The LUC:REN ratio represents *ProCYP94C1:LUC* activity relative to that of the internal control (*REN* driven by the 35S promoter). Data are mean ± SD, *n* = 3 repeats. Asterisks indicate significant*** differences between *35S:GFP* and *35S:EIL1-GFP* effector constructs. ( *P*<0.001; Student’s *t* test). **(D)** Schematic representation of *CYP94C1* showing the design of ChIP-qPCR and the EMSA. The vertical red line indicates the putative EBS; blue arrows indicate PCR amplicons detected by ChIP-qPCR; the short green line indicates the DNA probe used in the EMSA. **(E)** ChIP-qPCR results showing the enrichment of EIL1-GFP in the *CYP94C1* promoter. PCR amplicons are indicated in **(D)**. Tomato *ACTIN2* was used as a nonspecific target. Data are mean ± SD, *n* = 3 repeats. Asterisks indicate significant differences between MG and Br+4 fruits. (\*\*\**P*<0.001; Student’s *t* test). ns, not significant. **(F)** EMSA results showing that EIL1 directly binds to the *CYP94C1* promoter. A labeled probe incubated together with the GST protein was used as a negative control. Unlabeled probe (5-, 10-, and 20-fold excess) and mutant probe (20-fold excess) were used for competition.

The above findings, together with the fact that *CYP94C1* is a ripening-induced gene (Figures 2A and 2B), suggest that CYP94C1 terminates JA signaling in ripe fruits by converting JA-Ile to inactive 12-COOH-JA-Ile.

### EIL activates *CYP94C1* expression via promoter binding

We then explored how *CYP94C1* expression is activated by ripening. The results of a previous cleavage under targets and tagmentation assay (Deng et al., 2023) identified *CYP94C1* as an EIL-bound gene, suggesting that *CYP94C1* is a direct transcriptional target of EIL. To test this possibility, we examined the expression pattern of *CYP94C1* in *EIL-RNAi* fruits (Deng et al., 2023). RT-qPCR assays revealed that ripening-induced expression of *CYP94C1* was significantly lower in *EIL-RNAi* fruits than in WT fruits (Figure 3A), indicating that EIL positively regulates *CYP94C1* expression. We then used a well-established dual-luciferase (LUC) reporter system (Hellens et al., 2005) to verify the positive effect of EIL on *CYP94C1* expression. For this purpose, we cloned a 4,880-bp *CYP94C1* promoter sequence and inserted it into a dual-LUC reporter vector to generate a *ProCYP94C1:LUC* reporter construct (Figure 3B). As expected, coexpression of *35S:EIL1-GFP* with *ProCYP94C1:LUC* in *Nicotiana benthamiana* leaves led to increased LUC activity (Figure 3C).

We then investigated whether EIL directly binds to the *CYP94C1* promoter. Chromatin immunoprecipitation quantitative PCR (ChIP-qPCR) analysis using *ProEIL1:EIL1-GFP* fruits (Deng et al., 2023) and an anti-GFP antibody revealed ripening-induced enrichment of EIL-GFP at the EIN3-binding site (EBS) at the *CYP94c1* promoter (Figures 3D and 3E). Furthermore, electrophoretic mobility shift assays (EMSAs) revealed that a recombinant EIL1-glutathione S-transferase (GST) fusion protein (EIL-GST), but not GST itself, directly bound to a DNA probe containing the EBS motif, and that this binding was outcompeted by the unlabeled probe but not by a probe containing a mutated EBS motif (Figure 3F). Taken together, our results demonstrate that EIL1 specifically binds to the EBS motif of *CYP94C1* and activates its expression.

### CYP94B1 and CYP94B2 attenuate JA signaling in leaves by converting JA-Ile to its less active form 12-OH-JA-Ile

Next, to understand the role of *CYP94B* genes in JA signaling, we investigated the enzyme activity of CYP94B2 only, because of the high sequence similarity between CYP94B1 and CYP94B2 (Supplemental Figures 2A and 2B). LC-MS/MS analysis detected high levels of 12-OH-JA-Ile and relatively low levels of 12-COOH-JA-Ile after incubation of JA-Ile with the liquid culture of yeast cells expressing CYP94B2 (Figures 2A–2C), indicating that CYP94B2 has the same type of enzyme activity as its *Arabidopsis* orthologs and mainly converts JA-Ile to its less active form, 12-OH-JA-Ile.

To further determine the function of *CYP94B* genes in JA-mediated defense response, we generated CRISPR/Cas9-mediated *CYP94B1* and *CYP94B2* knockout mutants. Two resulting double mutants, *cyp94b1-1 cyp94b2-1* and *cyp94b1-2 cyp94b2-2*, were selected for further analysis (Supplemental Figures 4A and 4B). Infection with *B. cinerea* led to significantly smaller necrotic lesions on the leaves of *cyp94b1 cyp94b2* double mutants than those on WT plants (Figures 4A and 4B). Consistently, fungal growth on double mutant leaves was much lower than that on WT leaves (Figure 4C). These results demonstrated that CYP94B1 and CYP94B2 negatively regulate leaf resistance to *B. cinerea*.

**Figure 4.**
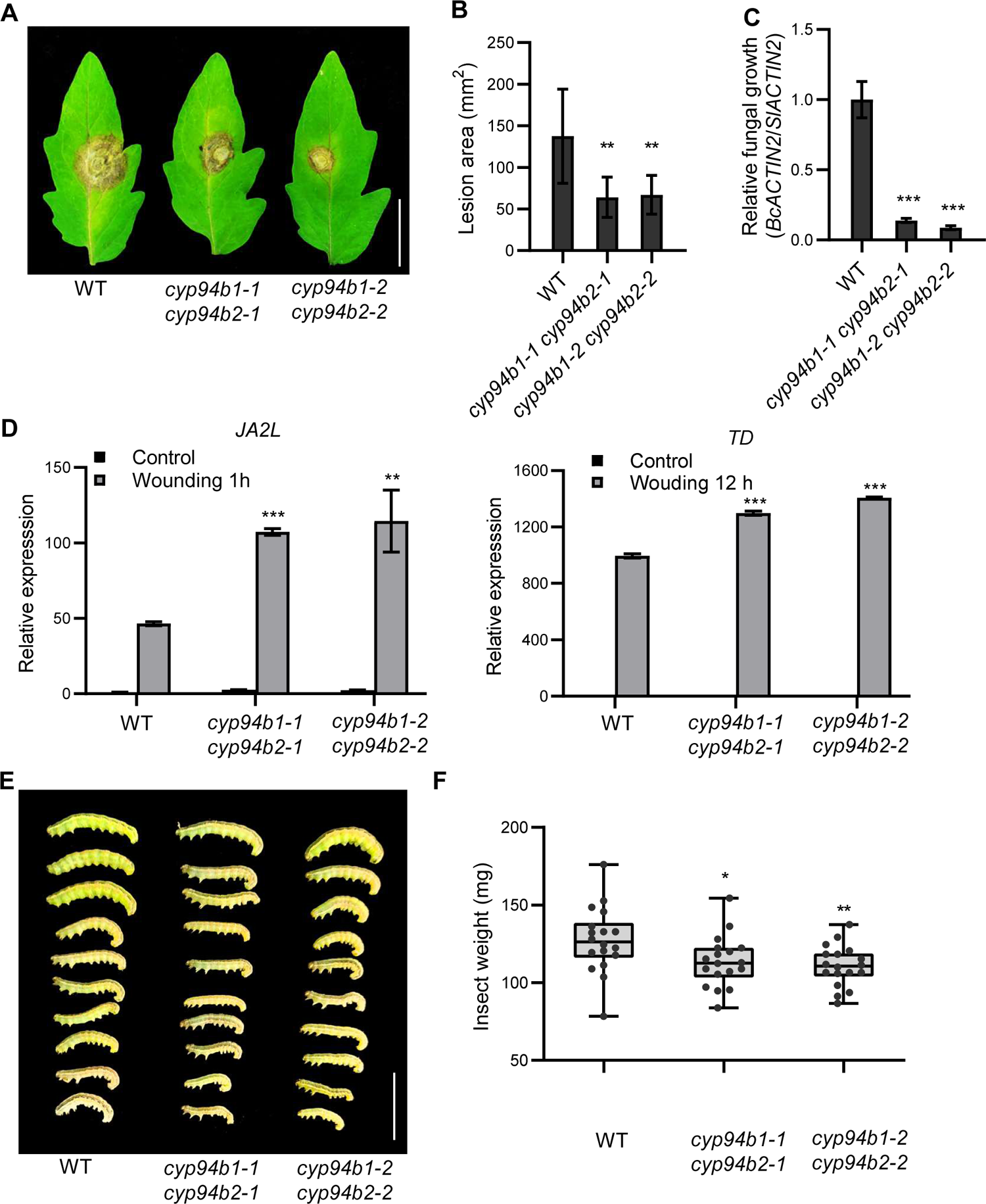
*CYP94B* negatively regulates JA-mediated defense response in leaves. **(A)** Representative images of *B. cinerea*-inoculated WT and *cyp94b1 cyp94b2* leaves. The images were taken at 3 DPI. Bar = 1.5 cm. **(B)** Lesion area on WT and *cyp94b1 cyp94b2* leaves. Data are mean ± SD, *n* = 8 repeats. **(C)** Quantification of fungal growth on WT and *cyp94b1 cyp94b2* leaves. Data are mean ± SD, *n* = 3 repeats. **(D)** RT-qPCR results showing the expression of *JA2L* and *TD* in WT and *cyp94b1 cyp94b2* plants in response to mechanical wounding. Data are mean ± SD, *n* = 3 repeats. **(E and F)** Representative image **(E)** and average weight **(F)** of larvae recovered at the end of the 6-day feeding trial using whole plants of the WT and c*yp94b1 cyp94b2* double mutants. Bar = 2 cm. Data are mean ± SD, *n* = 18 repeats. For **(B, C, D, and F)**, asterisks indicate significant differences between WT and *cyp94b1 cyp94b2* mutants (\**P*<0.05, \*\**P*<0.01, \*\*\**P*<0.001; Student’s *t* test).

We then investigated whether CYP94B genes is involved in defense responses to mechanical wounding. For this purpose, the *cyp94b1 cyp94b2* and WT seedlings were subjected to a standard wounding assay. RT-qPCR analysis revealed that wound-induced expression of *JA2L*, a well-characterized early JA-responsive gene targeted by MYC2 (Du et al., 2014; Du et al., 2017), was significantly higher in *cyp94b1 cyp94b2* double mutants than in WT plants (Figure 4D). Similarly, wound-induced expression of the late JA-responsive gene *TD* (Chen et al., 2005; Du et al., 2017) was also significantly higher in the double mutants than in the WT (Figure 4E). These results demonstrated that CYP94B1 and CYP94B2 negatively regulate defense responses to mechanical wounding.

Furthermore, we examined the performance of *cyp94b1 cyp94b2* double mutant plants to *Spodoptera exigua*, a globally significant agricultural pest. The average weight of larvae fed with *cyp94b1 cyp94b2* leaves was significantly lower than that of larvae fed with WT leaves (Figure 4F), indicating that CYP94B1 and CYP94B2 negatively regulate plant resistance to chewing insects.

These findings, together with the fact that *CYP94B1* and *CYP94B2* are wound-induced genes (Figures 2A and 2B), support the notion that CYP94B1 and CYP94B2 attenuate JA signaling in wounded leaves by converting JA-Ile to its less active form 12-OH-JA-Ile.

### MYC2 activates *CYP94B1* and *CYP94B2* via promoter binding

Next, we investigated how the expression of *CYP94B1* and *CYP94B2* is activated by wounding. As shown in Figure 1E, *CYP94B1* and *CYP94B2* expression increased rapidly and peaked 1 h after wounding, suggesting that they are early JA-responsive genes. Considering that MYC2 directly regulates the transcription of many early JA-responsive genes (Du et al., 2017), we speculated that C*YP94B1* and *CYP94B2* would be direct targets of MYC2. To test this hypothesis, we generated CRISPR/Cas9-mediated null mutants of *MYC2*. Two of the resulting mutants (*myc2-1* and *myc2-2*), each carrying a frameshift mutation resulting in premature translation termination (Supplemental Figure 5A), were selected and subjected to a standard wounding assay. RT-qPCR assays revealed that wound-induced *CYP94B1* and *CYP94B2* expression was dramatically lower in *myc2* mutants than in the WT (Figure 5A), indicating that MYC2 positively regulates the expression of *CYP94B1* and *CYP94B2*.

**Figure 5.**
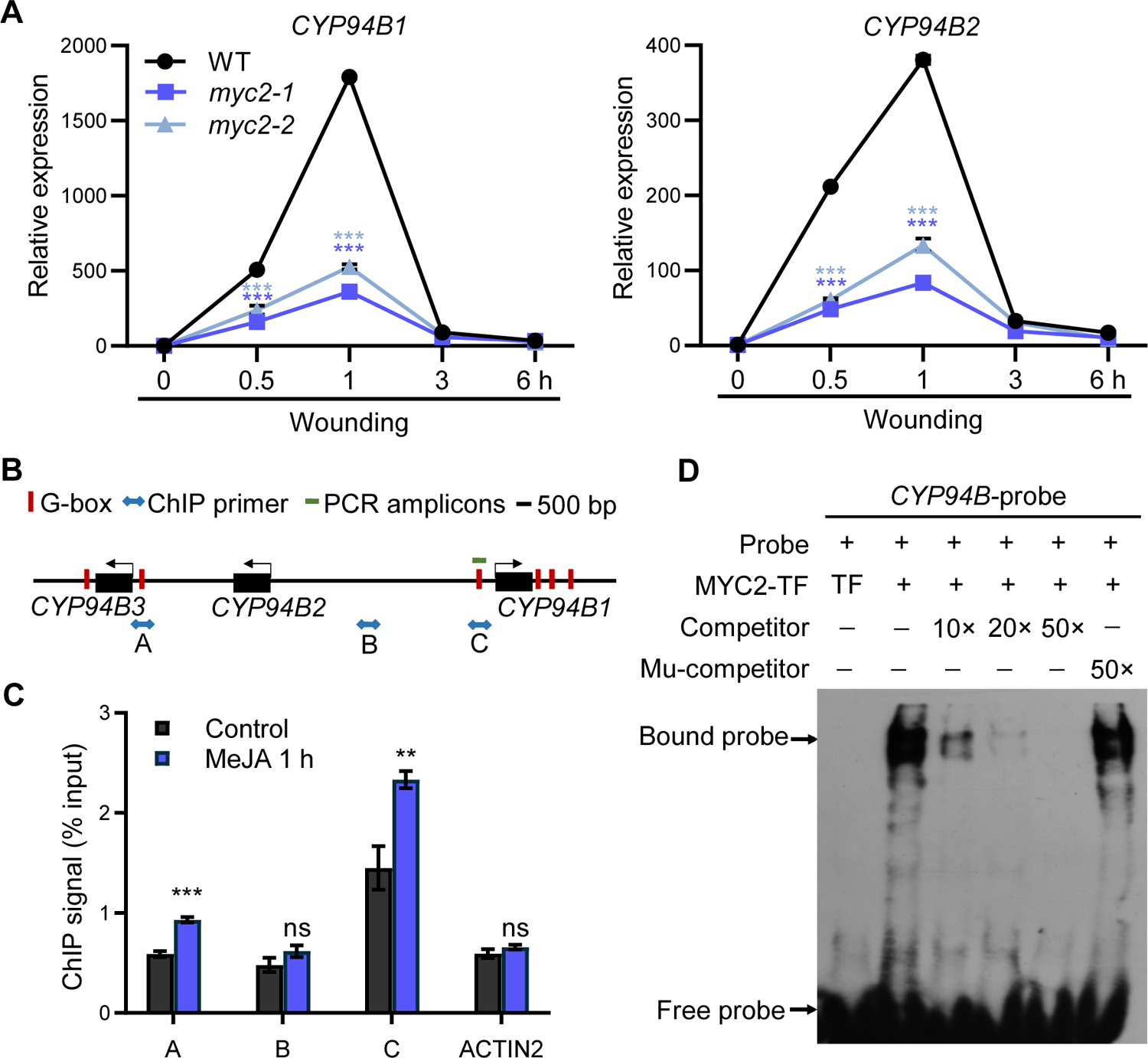
MYC2 regulates the expression of *CYP94B1* and *CYP94B2* by binding to their promoters. **(A)** RT-qPCR results showing wound-induced expression of *CYP94B1* and *CYP94B2* in WT and *myc2* leaves. Data are mean ± SD, *n* = 3 repeats. Asterisks indicate significant differences between WT and *myc2* mutants (\*\*\**P*<0.001; Student’s t test). **(B)** Schematic representation of the *CYP94B* gene cluster showing the design of ChIP-qPCR and the EMSA. Vertical red lines indicate putative EBSs; blue arrows indicate PCR amplicons detected by ChIP-qPCR; the short green line indicates DNA probe used in EMSA. **(C)** ChIP-qPCR results showing the enrichment of MYC2-GFP in the *CYP94B* promoter. PCR amplicons are indicated in **(B)**. Tomato *ACTIN2* was used as a nonspecific target. The fold enrichment of MYC2-GFP at the promoter was calculated against that of the *ACTIN2* promoter. Data are mean ± SD, *n* = 3 repeats. Asterisks indicate significant differences between MeJA-treated and untreated (control) plants. (\*\**P*<0.01, \*\*\**P*<0.001; Student’s *t* test). ns, not significant **(D)** EMSA results showing that MYC2 directly binds to the *CYP94B* promoter. A labeled probe incubated together with the TF protein was used as a negative control. Unlabeled probe (10-, 20-, and 50-fold excess) and mutant probe (50-fold excess) were used for competition.

We then investigated whether MYC2 regulates the expression of *CYP94B1* and *CYP94B2* by binding to their promoter regions. For this purpose, we generated *ProMYC2:MYC2-GFP* transgenic plants (Supplemental Figure 5B). ChIP-qPCR analysis using *ProMYC2:MYC2-GFP* plants and an anti-GFP antibody revealed JA-induced enrichment of MYC2-GFP on the G-box motifs of the *CYP94B* gene cluster (Figures 5B and 5C). Moreover, EMSAs showed that recombinant MYC2-trigger factor (TF) fusion protein (MYC2-TF), but not TF itself, bound with high affinity to the G-box motif in the *CYP94B1* promoter (Figure 5D). Together, our results demonstrate that EIL activates the expression of *CYP94B1* and *CYP94B2* by binding to the promoter regions of these genes.

### Editing of *CYP94* genes improves plant resistance to *B. cinerea* in fruits and leaves

Based on our findings, we propose a model to explain the action of different members of the *CYP94* gene family in orchestrating JA-mediated defense responses in tomato (Figure 6A). In wounded leaves, elevated JA-Ile levels lead to the de-repression of MYC2 and consequently to the activation of JA-mediated defense responses. Simultaneously, MYC2 directly activates the expression of *CYP94B1* and *CYP94B2*, the protein products of which in turn convert JA-Ile to its less active form 12-OH-JA-Ile, thereby attenuating JA signaling (Figure 6A). In ripening fruits, an ET burst at the onset of ripening leads to stabilization of EIL proteins, after which EIL orchestrates a hierarchical transcriptional cascade to promote the formation of fruit quality. Meanwhile, EIL directly activates the expression of *CYP94C1*, the protein product of which in turn converts JA-Ile to its inactive form 12-COOH-JA-Ile, thereby terminating JA signaling (Figure 6A).

**Figure 6.**
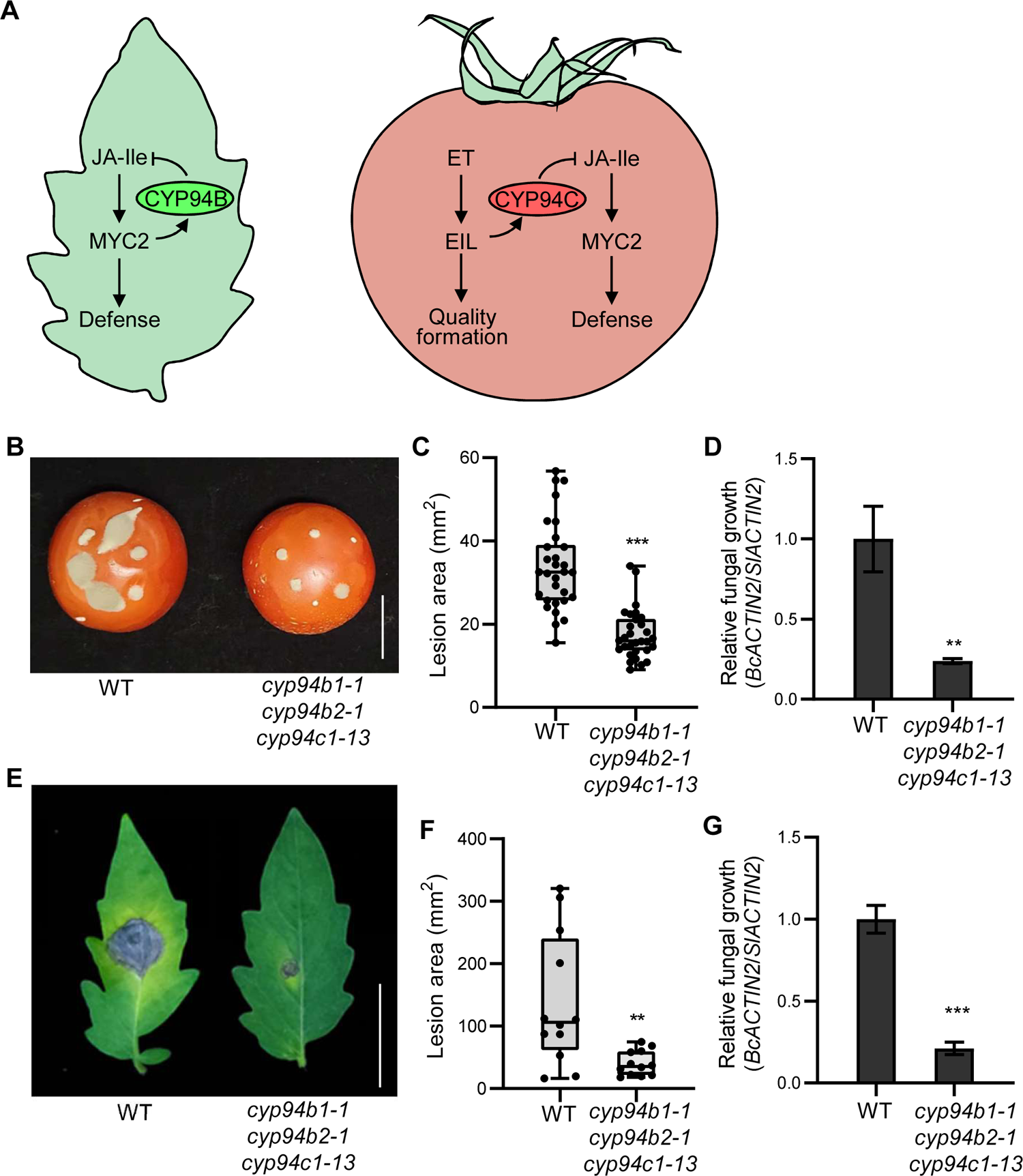
Simultaneous mutation of *CYP94B1*, *CYP94B2*, and *CYP94C1* leads to increased resistance against *B. cinerea*e in fruits and leaves. **(A)** A proposed working model showing the role the *CYP94* gene family members in orchestrating the JA-mediated defense response in tomato. **(B)** Representative images of *B. cinerea*-inoculated WT and *cyp94b1 cyp94b2 cyp94c1* fruits. The images were taken at 3 DPI. Bar = 2 cm. **(C)** Lesion area on WT and *cyp94b1 cyp94b2 cyp94c1* fruits. Data are mean ± SD, *n* = 30 repeats. **(D)** Quantification of fungal growth on WT and *cyp94b1 cyp94b2 cyp94c1* fruits. Data are mean ± SD, *n* = 3 repeats. **(E)** Representative images of B. *cinerea*-inoculated WT and *cyp94b1 cyp94b2 cyp94c1* leaves. The images were taken at 3 DPI. Bar = 2 cm. **(F)** Lesion area on WT and *cyp94b1 cyp94b2 cyp94c1* leaves. Data are mean ± SD, *n* = 12 repeats. **(G)** Quantification of fungal growth on WT and *cyp94b1 cyp94b2 cyp94c1* leaves. Data are mean ± SD, *n* = 3 repeats. For **(C, D, F, and G)**, asterisks indicate significant differences between WT and *cyp94b1 cyp94b2 cyp94c1* mutant (\*\**P*<0.01, \*\*\**P*<0.001; Student’s *t* test).

If our model is valid, simultaneous knockout of *CYP94B1*, *CYP94B2*, and *CYP94C1* should enhance JA-mediated defense in both fruits and leaves. To test this, we generated a *cyp94b1 cyp94b2 cyp94c1* triple mutant by crossing *cyp94b1-1 cyp94b2-1* mutant plants with the *cyp94c1-13* mutant. As expected, the fruits (Figures 6B–6D) and leaves (Figures 6E–6G) of the triple mutant were more resistant to *B. cinerea* than those of the WT. These results suggested that *CYP94* genes could be potentially be targeted to improve tomato resistance to necrotrophs.

## DISCUSSION

Fruit ripening leads to a wide range of changes in color, texture, flavor, aroma, and other quality attributes (Forlani et al., 2019; Klee and Giovannoni, 2011). Increased susceptibility to postharvest pathogens, especially necrotrophic pathogens, occurs alongside these changes. Although ripening-associated susceptibility to pathogens is widespread across different fruit species (Alkan and Fortes, 2015; Balsells-Llaurado et al., 2020; Blanco-Ulate et al., 2016; Cantu et al., 2009; Forlani et al., 2019; Haile et al., 2019; Li et al., 2022; Petrasch et al., 2019a; Petrasch et al., 2019b; Silva et al., 2023; Silva et al., 2021), the underlying mechanisms are still unclear. ET is a key signal governing the ripening in climacteric fruits, while JA is a major defense hormone that regulates plant resistance to mechanical wounding, chewing insects, and necrotrophic pathogens. Here, we report that ET-mediated ripening suppresses JA-mediated defense by activating the metabolism of JA-Ile, the most bioactive form of JA. Our study not only provides new insights into the mechanism responsible for the susceptibility of ripe fruits to necrotrophs but also offers a potential target for improving tomato fruit resistance to necrotrophs without compromising fruit quality.

### MYC2 and CYP94B form an autoregulatory feedback loop to attenuate JA signaling in green tissues

Because sustained activation of JA-mediated defense responses is detrimental, turning off JA signaling is as important as turning it on (Breeze, 2019; Liu et al., 2019; Zhai et al., 2020). We provide evidence that MYC2, together with its transcriptional targets *CYP94B1* and *CYP94B2*, forms transcriptional modules to attenuate JA signaling in leaves. First, wound-induced expression of *CYP94B1* and *CYP94B2* was impaired in *myc2* mutant plants. Second, ChIP-qPCR assays and EMSAs indicated that MYC2 directly bound the *CYP94B* promoters. Third, CYP94B2 converted JA-Ile to its less active form 12-OH-JA-Ile *in vitro*. Finally, *cyp94b1 cyp94b2* double mutants showed increased resistance to mechanical wounding, chewing insects, and *B. cinerea* infection. Thus, in addition to initiating and amplifying the transcriptional output of JA signaling (Chen et al., 2012; Du et al., 2017; Zhai et al., 2020), MYC2 forms an autoregulatory negative feedback circuit with CYP94B to attenuate JA signaling. Notably, the activation of *CYP94B* genes is already pre-programmed during the initiation phase of JA signaling. Therefore, CYP94B-mediated JA-Ile deactivation is an integral part of JA signaling. Considering that JA-Ile level and JA-mediated defense responses were high in unripe fruits, it is reasonable to speculate that the mechanism we described here also operates in unripe fruits.

In addition to the deactivation of JA-Ile, plants have evolved other autoregulatory feedback loops that ensure the proper attenuation or termination of JA signaling. For example, the activation of MYC2 induces the expression of *JAZ* repressors. Most *JAZ* genes contain a highly conserved JA-associated (Jas) intron, and alternative splicing involving the Jas intron generates a repertoire of dominant JAZ splice variants that are unable to interact with the receptor for degradation but still able to repress MYC2 (Chung et al., 2010; Chung and Howe, 2009; Moreno et al., 2013; Wu et al., 2020; Zhang et al., 2017). Thus, the generation of JAZ splice variants provides an efficient route for desensitizing JA signaling. Furthermore, MYC2 directly activates a small group of JA-inducible bHLH transcription factors, termed MYC2-TARGETED BHLH1 (MTB1), MTB2, and MTB3, which negatively regulate JA responses (Liu et al., 2019). MTB proteins disrupt the interaction of MYC2 with its coactivator and impair the binding of MYC2 to its target promoters, therefore contributing to the termination of JA signaling (Liu et al., 2019). In this context, our findings stress again the role of MYC2 as a ‘master switch’ in JA signaling.

### EIL relays the ET signal to CYP94C1 to terminate JA signaling during fruit ripening

We also identified the interaction node between ET-mediated ripening and JA-mediated defense. We provide evidence that EIL, together with its direct target *CYP94C1*, forms a transcriptional module to terminate JA signaling during fruit ripening. First, the ripening-induced decrease in the JA-Ile level was strongly dependent on EIL function. Second, ripening-induced expression of *CYP94C1* was much lower in *EIL-RNAi* fruits than in WT fruits. Third, the results of ChIP-qPCR and EMSAs showed that EIL directly binds to the *CYP94C1* promoter. Fourth, CYP94C1 converted bioactive JA-Ile to its inactive form 12-COOH-JA-Ile *in vitro*. Finally, knockout of *CYP94C1* increased ripe fruit resistance to *B. cinerea*. These results support a scenario in which EIL relays ET signals to *CYP94C1* to terminate JA signaling during fruit ripening, thereby resulting in increased fruit susceptibility to necrotrophs. Thus, CYP94C1-mediated JA-Ile deactivation is an integral part of ET-mediated ripening.

This finding, together with those of previous studies (Deng et al., 2023; Huang et al., 2022), suggest that ET has a dual role in regulating fruit ripening. On the one hand, ET promotes fruit softening and accumulation of flavor and aroma compounds, which make fruits more attractive to and edible by animals. On the other hand, it alleviates JA-mediated defense responses, rendering fruits more sensitive to necrotroph infection. Considering that fruits protect the developing seeds and serve as the vehicle for seed dispersal (Fenn and Giovannoni, 2021; Forlani et al., 2019; Tanksley, 2004), it is likely that both of these effects of ET facilitate the dispersal of mature seeds in nature.

It has been recently reported that *B. cinerea* infection accelerates ripening processes, such as cell wall disassembly, to facilitate its colonization (Silva et al., 2023). ET production rapidly increases in *B. cinerea*-inoculated fruits (Silva et al., 2023). Considering the crucial role of ET in tomato fruit ripening (Deng et al., 2023; Huang et al., 2022), it is reasonable to speculate that *B. cinerea* hijacks the ET pathway to promote its infection. In this context, it would be interesting to examine whether the EIL-CYP94C1 module is hijacked by *B. cinerea* as an infection strategy.

In *Arabidopsis*, all three major *CYP94* genes (*CYP94B1*, *CYP94B3*, and *CYP94C1*) are strongly induced by wounding (Heitz et al., 2012; Koo et al., 2011). Although tomato *CYP94B1* and *CYP94B2* show similar wounded-induced expression patterns to those of their *Arabidopsis* orthologs, *CYP94C1* expression is specifically induced by ripening but not by wounding. In addition, our results revealed that the functions of CYP94B2 and CYP94C1 in JA-Ile turnover are different. While CYP94B2 primarily converts JA-Ile to its less active form 12-OH-JA-Ile, CYP94C1 tends to convert JA-Ile to its inactive form 12-COOH-JA-Ile. Taking these data into consideration, it is plausible that differences in the expression and enzymatic activities of *CYP94* gene family members provide precise control over JA-mediated defense responses in tomato.

### *CYP94* gene family members are potential targets for improving tomato resistance to necrotrophs

As most fruit quality attributes are determined during ripening, breeding fruit resistance to necrotrophs without compromising the quality of ripened fruit is a big challenge for many crops. For example, although the natural mutation *nor* results in robust resistance to *B. cinerea* (Cantu et al., 2009), it adversely affects nutritional and sensory quality attributes (Adaskaveg et al., 2021; Garg et al., 2008; Wang et al., 2020). Our results revealed that CRISPR/Cas9-mediated mutation of *CYP94C1* led to increased resistance to *B. cinerea* without affecting the ripening process. From this perspective, we suggest that *CYP94C1* should be added to the list of other genes that have been reported as potential targets for the enhancement of tomato fruit resistance to necrotrophs, such as *PL* (Silva et al., 2021) and the combination of *POLYGALACTURONASE 2a* (*PG2a*) and *EXPANSIN 1* (*EXP1*) (Cantu et al., 2008). Furthermore, simultaneous mutation of *CYP94B1*, *CYP94B2*, and *CYP94C1* enhanced the JA-mediated defense response in fruits and leaves, suggesting that targeting *CYP94* genes could lead to the development of new strategies for crop protection. In future studies, it will be critical to determine whether the disruption of these genes affects important agronomic tomato traits, such as yield and fruit quality. Another interesting direction for future exploration is to test whether the functions of *CYP94C* and *CYP94B* are conserved in different fruit species.

## METHODS

### Plant materials and growth conditions

Seeds of the tomato cultivar Ailsa Craig (AC; LA2838A) were obtained from the Tomato Genetics Resource Center (https://tgrc.ucdavis.edu) and used as the WT in this study. *EIL-RNAi* and *ProEILpro:EIL1-GFP* (Deng et al., 2023) seeds were obtained from our own stocks. Tomato seeds were germinated on moistened filter paper at room temperature and then sown in 32-cell plastic flats. Seedlings were grown in a growth chamber under controlled conditions (16 h of light [200 µE m^-2^ s^-1^] at 26°C and 8 h of dark at 18°C with 60% relative humidity). Thirty-d-old seedlings were transplanted into pots with soil and grown in a naturally illuminated greenhouse (16 h day [26 ± 2°C]/8 h night [20 ± 2°C] cycle and 60% relative humidity). Flowers were tagged at anthesis. Fruits were harvested at 35, 42, and 46 days post anthesis (DPA), which corresponds to the MG stage, Br stage, and Br+4 stage of the WT, respectively. For RT-qPCR, the pericarps of five fruits were pooled for each sample. *N. benthamiana* seeds were obtained from our own stocks and directly sown in soil in 8-cm diameter pots for 10 d before being transplanted to the 32-cell plastic flats. *N. benthamiana* plants were grown under the same conditions as the tomato seedlings.

### Plasmid constructs and plant transformation

To generate the *ProMYC2:MYC2-GFP* construct, the 35S promoter of the *pCAMBIA1300-GFP* vector was replaced by a DNA fragment containing the 1,796-bp *MYC2* native promoter and the full-length coding sequence (CDS) of *MYC2* without stop codon using SalI (Thermo Fisher, FD0644) and KpnI (Thermo Fisher, FD0524) restriction endonucleases. The resulting construct was introduced into the tomato cultivar AC via *Agrobacterium tumefaciens* (LBA4404)-mediated transformation (Deng et al., 2018). Transformants were selected on the basis of their resistance to hygromycin B. Homozygous T2 transgenic plants were used for further experiments. The primers used for plasmid construction are listed in Supplemental Data Set 1.

### CRISPR/Cas9-mediated mutation

Null mutations of *CYP94C1*, *CYP94B1*, *CYP94B2*, and *MYC2* were generated using the tomato *U6* promoter-controlled CRISPR/Cas9 system as described previously (Yang et al., 2019). Briefly, two sets of primers containing guide RNA (gRNA) sequences of *CYP94C1* were used in PCR to generate a tomato *U6-26-CYP94C1-gRNA* cassette. The resulting *U6-26-CYP94C1-gRNA* cassette was then cloned into the binary vector *pTX041* (Deng et al., 2018) to form a *pTX041-CYP94C1* construct. *pTX041-CYP94B1&2* and *pTX041-MYC2* constructs were generated using the same protocol. The final constructs were then transformed into AC via *Agrobacterium tumefaciens* (LBA4404)-mediated transformation. The CRISPR/Cas9-mediated mutations were genotyped by PCR amplification and DNA sequencing. Homozygous T2 lines without Cas9 were identified for further experiments. The primers used for plasmid construction and genotyping are listed in Supplemental Data Set 1.

### Plant treatment and gene expression analysis

MeJA treatment was performed as described previously (Liu et al., 2019). Eighteen-d-old tomato seedlings (with two expanded leaves and a third emerging leaf) were wounded with a hemostat across the midrib of all leaflets of the lower and upper leaves. Plants with wounded leaves were incubated under continuous light. Whole leaves were harvested at each sampling time point and used for total RNA extraction. Leaves of unwounded plants were harvested as a control. The leaves of five plants were pooled for each sample. Total RNA was extracted from each sample using TRIzol reagent (Invitrogen, 15596018) according to the manufacturer’s instructions. The quality of the total RNA was determined using a NanoDrop spectrophotometer (Thermo Fisher). Each sample (2 μg) of total RNA was used to synthesize first-strand cDNA with a PrimeScript RT Kit with gDNA eraser (Takara, RR0447A). RT-qPCR was performed using a Roche LightCycler 480 system with a SYBR Fast qPCR Kit (KAPA Biosystems, KK4601). The expression levels of the target genes were normalized against that of tomato *ACTIN2*. The error bars represent the SDs of three biological replicates. The primers used to quantify gene expression levels are listed in Supplemental Data Set 1.

### Ethylene measurement

ET production was measured as described previously (Deng et al., 2023; Yin et al., 2015) with modification. For each sample, three fruits at the same stage were harvested and immediately enclosed in 500-mL air-tight glass containers for 1 h at 20°C. ET emission was determined by gas chromatography using a Shimadzu GC2014 equipped with a flame ionization detector. The error bars represent the SDs of three biological replicates.

### JA-Ile and α-tomatine measurement

For JA-Ile measurement, the pericarps of fruits at corresponding stages were harvested and immediately ground to a powder in liquid nitrogen. JA-Ile extraction and measurement were carried out as described previously (Fu et al., 2012; Liu et al., 2019), with minor modifications. Briefly, five volumes of ice-cold 90% methanol containing JA-d5 as an internal standard were added to one fresh weight (600 mg) of frozen powder in the tube. Homogenates were cleared by two successive centrifugations at 20,000 × g, and the supernatants were used for ultra-performance liquid chromatography-mass spectrometry analysis as described by Heitz et al. (2012). For α-tomatine measurement, approximately 300 mg of powder was extracted with MeOH (300 µL) containing 10 µg/mL genistein as an internal standard. The extraction and high performance liquid chromatography coupled to Fourier transform ion cyclotron resonance mass spectrometry analysis were performed as described previously (Iijima et al., 2013). The content of α-tomatine was calculated using the peak area ratio of tomatine and the internal control.

### *In vitro* enzyme activity assays

*In vitro* enzyme activity assays were performed as described previously (Nomura et al., 2013; Pompon et al., 1996) with minor modifications. The CDSs of *CYP94B2* and *CYP94C1* were amplified from cDNA and cloned into the *pYeDP60* vector (Pompon et al., 1996). The resulting construct was transformed into yeast strain WAT11 engineered to co-produce *Arabidopsis* NADPH P450 reductase-1. The transformed colonies were inoculated into 10 mL of SGI medium (Pompon et al., 1996) and grown at 30°C for 1 d (200 rpm). Then, 1 mL SGI-cultured yeast medium was mixed with 10 mL SLI medium (Pompon et al., 1996) and incubated at 28°C overnight. The suspension was diluted with fresh SLI medium to an OD_600_ of 0.4. Five milliliter of the diluted suspension was incubated with 1 µg of JA-Ile substrate at 28°C for 16 h. The incubated suspensions were sonicated on ice (20 times for 5 s/5 s) and the pH was adjusted to 4.0 using concentrated HCl. The products were extracted with ethyl-acetate, and the acidic ethyl-acetate phase was evaporated until dry. The residual solid was dissolved in methanol and analyzed by LC-MS/MS. The MassLynx program was used for data acquisition and analysis. The transitions were, in negative mode, JA-Ile 322>130, 12-OH-JA-Ile 338>130, and 12-COOH-JA-Ile 352>130 and matched that of a corresponding standard.

### *B. cinerea* inoculation assays

*B. cinerea* inoculation assays were performed as described previously (Du et al., 2017). Briefly, *B. cinerea* isolate B05.10 was grown on V8 agar (34% V8 original juice, 0.2% CaCO3, and 2% agar) for 14 d at 22°C in the dark. Spore suspensions were prepared by harvesting the spores in 1% Sabouraud Maltose Broth (4% maltose and 1% peptone), filtering them through nylon mesh to remove hyphae, and adjusting the concentration to 1×10^5^ spores/mL. For the pathogenicity test, 4-week-old tomato leaves were picked and placed in Petri dishes containing 8% agar medium, and then 10 µL of spore suspension was inoculated. For fruits, the surface was punctured to a depth of 2 mm depth and a diameter of 1 mm, and then 10 µl of spore suspension was inoculated. The lesion area was measured at 3 DPI using ImageJ. To quantify *B. cinerea* growth, total DNA was extracted from infected leaves or fruits using a DNeasy Plant Mini Kit (Qiagen, 74904). The fungal growth was determined using qPCR amplification of *B. cinerea ACTIN2* gDNA relative to that of tomato *ACTIN2* gDNA, as described previously (Wang et al., 2017). The primers used for qPCR are listed in Supplemental Data Set 1.

### Insect feeding trials

Insect feeding assays were performed as described previously (You et al., 2019), with minor modifications. Briefly, *S. exigua* eggs were hatched at 26°C. Third-instar *S. exigua* larvae were transferred to a Petri dish and starved for 12 h before being used in the feeding experiment. For each genotype, more than twenty third-instar *S. exigua* larvae of a similar weight were placed in 150-mm plastic Petri dishes with leaflets from 4-week-old plants. The leaves in each Petri dish were replaced every 2 days. Larvae weight was measured 6 days later.

### Transient expression assays in *N. benthamiana* leaves

Transient transcriptional activity assays were performed using *N. benthamiana* leaves, as described previously (Sun et al., 2020). The 4880-bp promoter sequence of *CYP94C1* was amplified from AC genomic DNA and cloned into the *pGreenII 0800-LUC* vector (Hellens et al., 2005) for use as a reporter. The Renilla luciferase (REN) gene under the control of the cauliflower 35S promoter in the *pGreenII 0800-LUC* vector was used as an internal control. The *EIL1* CDS was cloned into the pGWB5 vector under the control of the 35S promoter and was used as an effector. The primers used for plasmid construction are listed in Supplemental Data Set 1. Firefly LUC and REN activities were measured using the Dual-Luciferase Reporter Assay System (Promega, E1910) following the manufacturer’s instructions, and the LUC:REN ratio was calculated and presented. Data from three independent biological replicates were collected, and error bars represent the SDs of three biological replicates.

### ChIP-qPCR assay

ChIP-qPCR assays were performed as described previously (Deng et al., 2023), with slight modifications. In short, the pericarps of *ProEIL1:EIL1-GFP* fruits harvested at 35, 42, and 46 DPA, and the leaves of *ProMYC2:MYC2-GFP* were crosslinked in 1% formaldehyde under a vacuum for 5 min. The crosslinking reaction was stopped by adding 0.125 M glycine, and the vacuum was extended for 5 min. The crosslinked samples were ground in liquid nitrogen. The chromatin complex was isolated, resuspended in lysis buffer (50 mM HEPES [pH 7.5], 150 mM NaCl, 1 mM EDTA, 1% SDS, 1% Triton X-100, 0.1% sodium deoxycholate, 1 mM phenylmethylsulfonyl fluoride, and 1× Roche Protease Inhibitor Cocktail) and sonicated to reduce the average DNA fragment size to ∼500 bp. Then, 50 μL of the sheared chromatin was removed and saved as an input control. The remaining chromatin was incubated with anti-GFP antibody (Abcam, ab290) at 4°C overnight. The immunoprecipitated chromatin–protein complex was sequentially washed with low-salt washing buffer (20 mM Tris-HCl [pH 8.0], 150 mM NaCl, 2 mM EDTA, 0.2% [w/v] SDS, and 0.5% [v/v] Triton X-100), high salt washing buffer (20 mM Tris-HCl [pH 8.0], 500 mM NaCl, 2 mM EDTA, 0.2% [w/v] SDS, and 0.5% [v/v] Triton X-100), LiCl washing buffer (10 mM Tris-HCl [pH 8.0], 25 mM LiCl, 1 mM EDTA, 0.5% [w/v] Nonidet P-40, and 0.5% [w/v] sodium deoxycholate), and TE buffer (10 mM Tris–HCl [pH 8.0] and 1 mM EDTA). The sample was then eluted with elution buffer (1% SDS and 100 mM NaHCO_3_). Reversal of protein-DNA crosslinking was performed by incubating the immunoprecipitated complex at 65°C overnight. The immunoprecipitated DNA was purified using AFTMag NGS DNA Clean Beads (ABclonal, RK20257). The ChIP signals were quantified using qPCR and displayed as the percentage of precipitated DNA relative to that of input DNA. The fold enrichment in selected regions was normalized against the nonspecific binding region of the tomato *ACTIN2* promoter. The error bars represent the SDs of three biological replicates. Each replicate was collected from five pooled fruits or seedlings at the same stage. The primers used for ChIP-qPCR are listed in Supplemental Data Set 1.

### EMSAs

EMSAs were carried out as described previously (Sun et al., 2020) with slight modification. EIL1-GST and GST proteins were synthesized using the TnT SP6 High-Yield Wheat Germ Protein Expression System (Promega, L3260), according to the manufacturer’s instructions. MYC2-TF and TF proteins were expressed in *Escherichia coli* BL21 (DE3) cells. Oligonucleotide probes were synthesized and labeled with biotin at the 5′ end by Invitrogen. The EMSA was performed using a Chemiluminescent EMSA kit (Beyotime, GS009). Biotin-labeled probes were incubated with proteins at room temperature for 10 min and free and bound probes were separated in an acrylamide gel. Unlabeled WT probes and mutated probes (containing altered EBS or G-box) were used as competitors. The probes used for the EMSA are listed in Supplemental Data Set 1.

### Sequence alignment and phylogenetic tree construction

Amino acid sequence alignment was preformed using Clustal X2 with default parameters (Larkin et al., 2007). Phylogenetic trees were constructed by MEGA version 7.0 (Kumar et al., 2016) using the neighbor-joining method. The branches were compared with bootstrap support values from 500 replicates for each node.

### Statistical analysis

A two-tailed Student’s *t* test was used to determine significance (\**P*<0.05; \*\**P*<0.01; \*\*\**P*<0.001; ns, not significant). A summary of the statistical analysis is shown in Supplemental Data Set 2.

### ACCESSION NUMBERS

Sequence data of tomato genes analyzed in this study are available in the SOL Genomics Network (http://solgenomics.net/) or National Center for Biotechnology Information (https://www.ncbi.nlm.nih.gov/) databases under the following accession numbers: *CYP94B1*, Solyc03g111300; *CYP94B2*, Solyc03g111290; *CYP94B3*, Solyc03g111280; *CYP94C1*, Solyc10g083400; *CYP94C2*, Solyc06g074420; *EIL1*, Solyc06g073720; *EIL2*, Solyc01g009170; *EIL3*, Solyc01g096810; *EIL4*, Solyc06g073730; *MYC2*, Solyc08g076930; *JA2L*, Solyc07g063410; *TD*, Solyc09g008670; *ACTIN2*, Solyc11g005330; and *BcACTIN2*, XM_024697950.1.

## Supporting information

Supplemental Figuers 1-5

Supplemental Data Sets 1-2

## SUPPLEMENTAL INFORMATION

Supplemental information is available at

## FUNDING

This study was supported by the National Natural Science Foundation of China (31991183, 32161133018, and U22A20459), the Strategic Priority Research Program of the Chinese Academy of Sciences (XDPB16), Hainan Yazhou Bay Seed Lab (B21HJ0109), and the Beijing Joint Research Program for Germplasm Innovation and New Variety Breeding (G20220628003).

## AUTHOR CONTRIBUTIONS

C.L. and L.D. conceived and designed the overall research. C.L. supervised the project. T.Y. and L.D. performed most of the experiments. Q.X. measured α-tomatine. P.X. and J.C. measured JA-Ile. C.S. and F.W. generated transgenic tomato plants. H.Z., T.H., and C.-B.L. helped grow the plants. C.L., L.D., T.Y., and M.A. wrote the manuscript.

## ACKNOWLEDGMENTS

We thank Prof. Xiaoya Chen (CAS Center for Excellence in Molecular Plant Science, Institute of Plant Physiology and Ecology, Chinese Academy of Sciences, Shanghai, China) for sharing the WAT11 yeast strain and the pYeDP60 vector. We thank Prof. Jinsong Zhang (Institute of Genetics and Developmental Biology, Chinese Academy of Sciences, Beijing, China) for helping with ET production measurement.

## Notes

### Competing Interest Statement

The authors have declared no competing interest.

