## Supplemental Figuers 1-5 for "Tomato CYP94C1 terminates jasmonate signaling during fruit ripening by inactivating bioactive jasmonoyl-L-isoleucine"

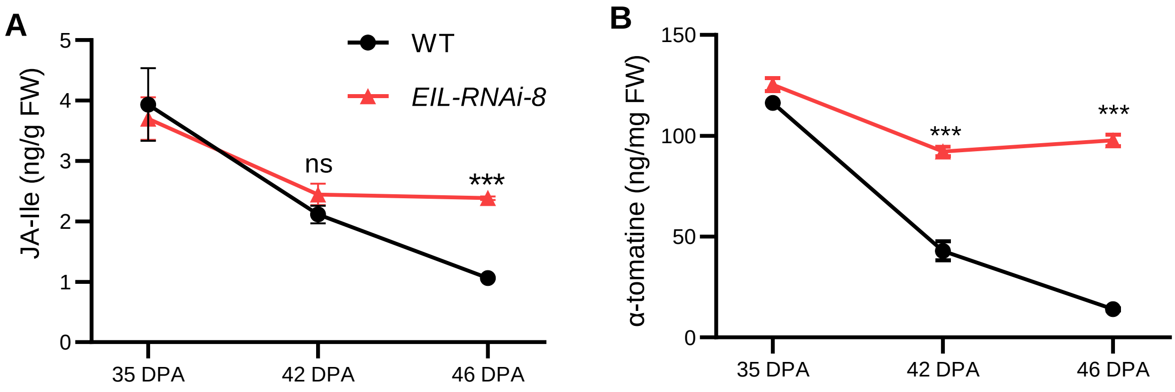


**Supplemental Figure 1. *EIL* knockdown impairs the ripening-induced decrease in JA-Ile and α-tomatine contents. Related to Figure 1.**

**(A and B)** JA-Ile **(A)** and α-tomatine **(B)** contents in WT and *EIL-RNAi* fruit harvested at the indicated stages. Fruits were harvested at 35, 42, and 46 DPA, which corresponds to the MG stage, Br stage, and Br+4 stage in the WT, respectively. Data are mean ± SD, n = *3* repeats. Asterisks indicate significant differences between WT and *EIL-RNAi* fruits (***$P < 0.001$; Student's *t* test). ns, not significant.

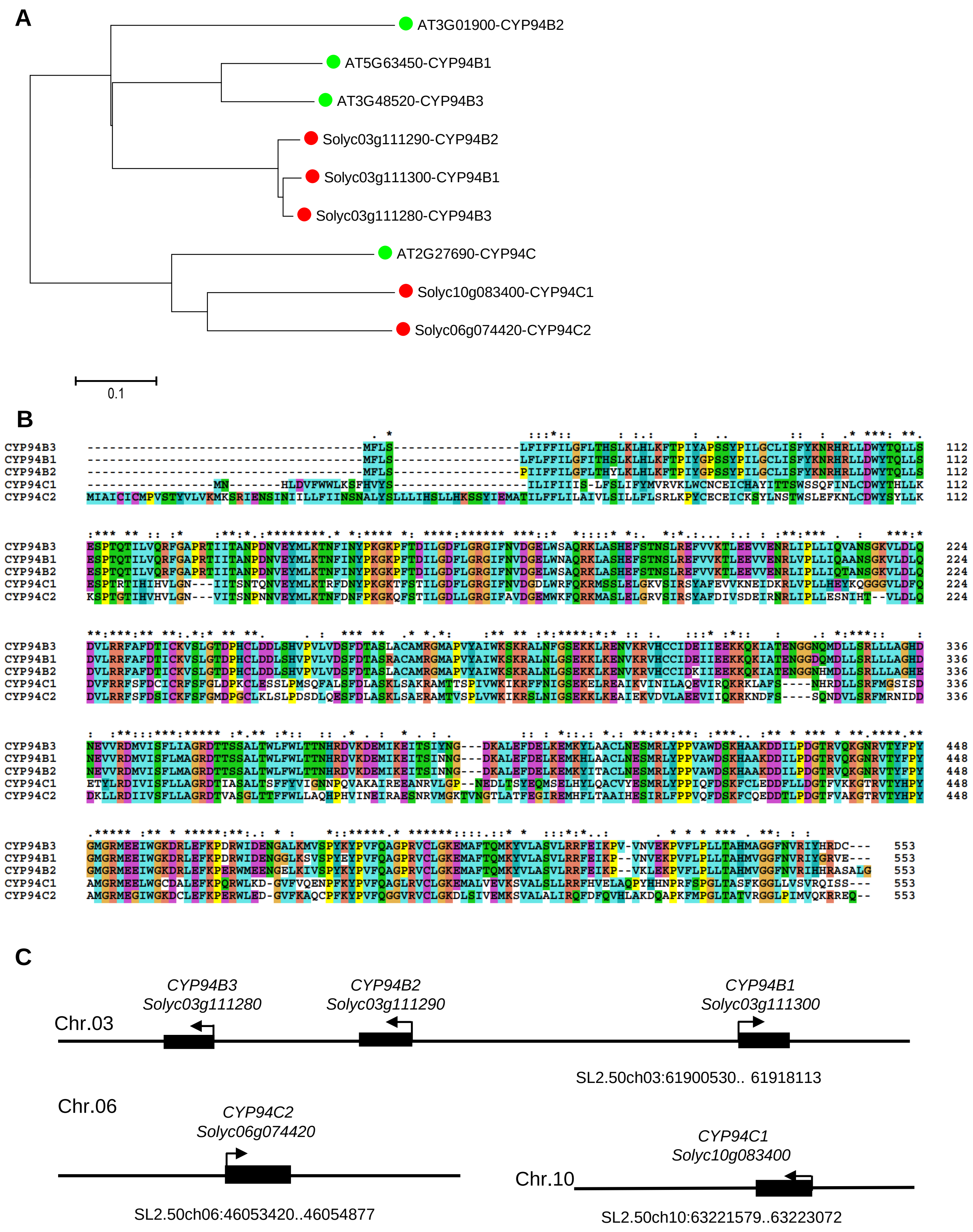


**Supplemental Figure 2. Sequence analysis of tomato CYP94 proteins and their *Arabidopsis* orthologs. Related to Figure 1.**

**(A)** Phylogenetic analysis of tomato CYP94 proteins (red dots) and their *Arabidopsis* orthologs (green dots). The phylogenetic tree was constructed by MEGA version 7.0 program using neighbor-joining method. The scale bar indicates the average number of amino acid substitutions per site.

**(B)** Amino acid sequence alignments of tomato CYP94B1, CYP94B2, CYP94B3, CYP94C1 and CYP94C2. The sequence alignment was preformed using Clustal X2 with default parameters.

**(C)** Chromosomal localization of tomato *CYP94* genes.

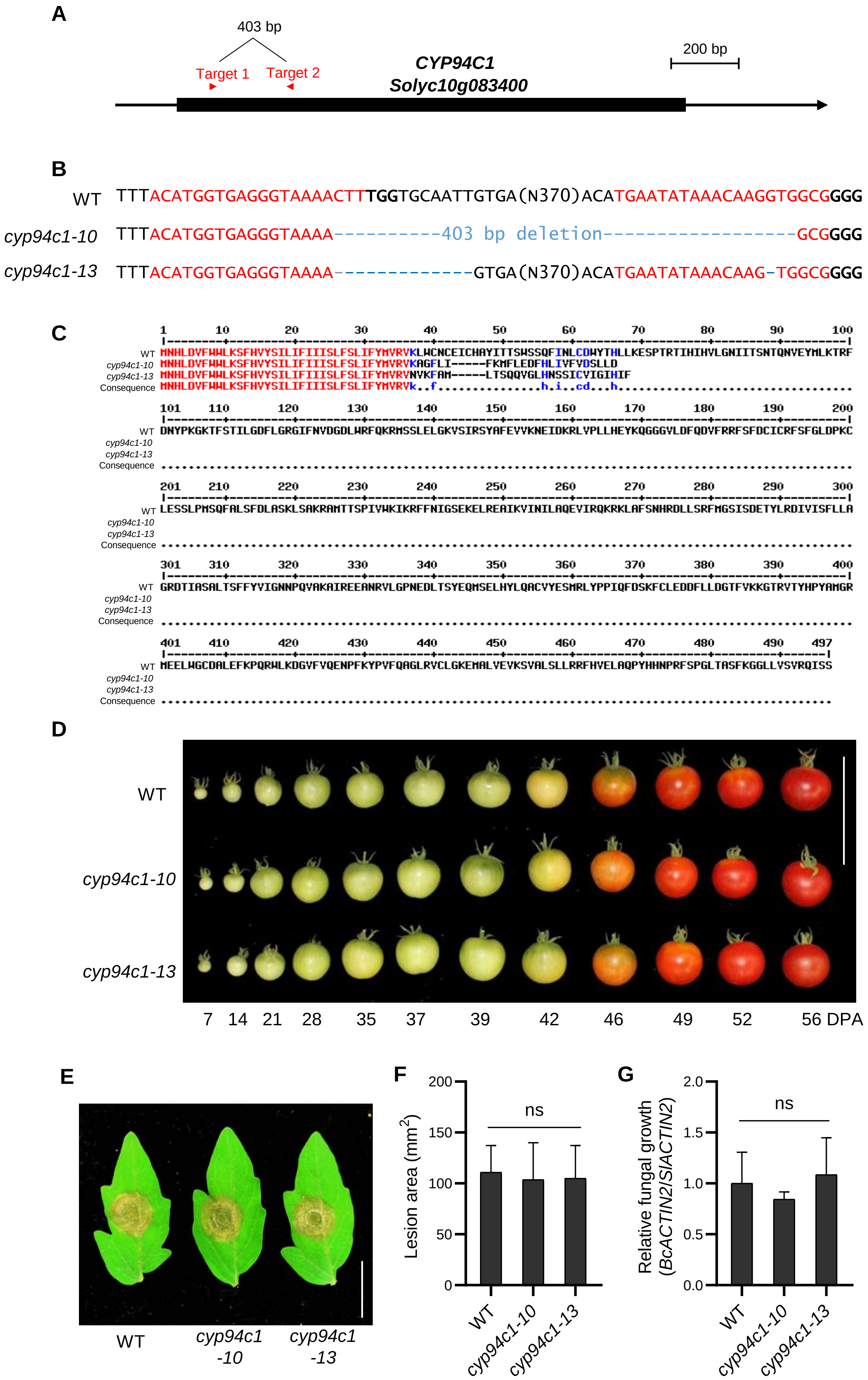
A
403 bp
Target 1
Target 2
CYP94C1
Solyc10g083400
200 bp
B
WT TTTACATGGTGAGGGTAAAACTTTGGTGCAATTGTGA(N370)ACATGAATATAAACAAGGTGGCGGGG
cyp94c1-10 TTTACATGGTGAGGGTAAAA----------403 bp deletion-----------------GCGGGG
cyp94c1-13 TTTACATGGTGAGGGTAAAA-------------GTGA(N370)ACATGAATATAAACAAG-TGGCGGGG
C
WT
cyp94c1-10
cyp94c1-13
Consequence
D
WT
cyp94c1-10
cyp94c1-13
7 14 21 28 35 37 39 42 46 49 52 56 DPA
E
WT
cyp94c1 -10
cyp94c1 -13
F
Lesion area (mm²)
ns
WT
cyp94c1-10
cyp94c1-13
G
Relative fungal growth (BcACTIN2/SlACTIN2)
ns
WT
cyp94c1-10
cyp94c1-13

**Supplemental Figure 3. Generation and phenotype of *cyp94c1* mutants. Related to Figure 2.**

**(A)** Schematic representation of *CYP94C1* showing the design of CRISPR/Cas9-mediated gene editing.

**(B)** DNA sequence analysis of *cyp94c1* null alleles generated by gene editing. The sgRNA targets and PAM are highlighted in red and bold font, respectively. The blue dashes and letters indicate deletions and insertions, respectively.

**(C)** Predicted amino acid sequences encoded by *cyp94c1* null alleles. The sequence alignment was preformed using MultAlin online program (http://multalin.toulouse.inra.fr/multalin/) with default parameters.

**(D)** Photographs of WT and *cyp94c1* fruits at the indicated stages. Bar = 10 cm.

**(E)** Representative images of *B. cinerea*-inoculated WT and *cyp94c1* leaves. The images were taken at 3 DPI. Bar = 2 cm.

**(F)** Lesion area on WT and *cyp94c1* leaves. Data are mean ± SD, $n$ = 7 repeats. ns, not significant.

**(G)** Quantification of fungal growth on WT and *cyp94c1* leaves. Data are mean ± SD, $n$ = 3 repeats. ns, not significant.

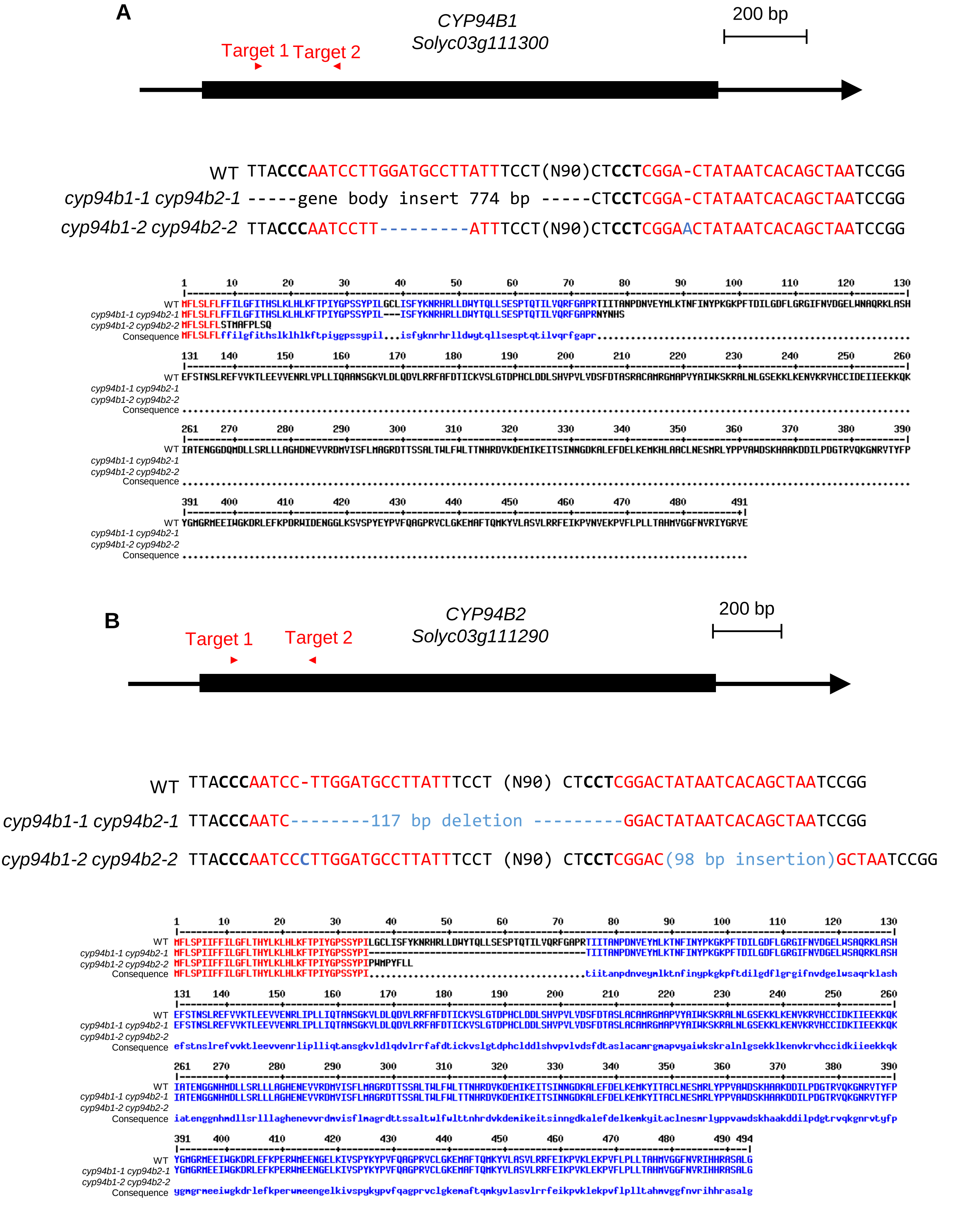


**Supplemental Figure 4. Generation of *cyp94b1 cyp94b2* double mutants. Related to Figure 4.**

**(A and B)** Sequence analysis of *CYP94B1* **(A)** and *CYP94B2* **(B)** in double mutants. Upper panel: schematic representation of *CYP94B1* or *CYP94B2* showing the design of CRISPR/Cas9-mediated gene editing. Middle panel: DNA sequence analysis of null alleles generated by gene editing. Bottom panel: predicted amino acid sequences encoded by null alleles.

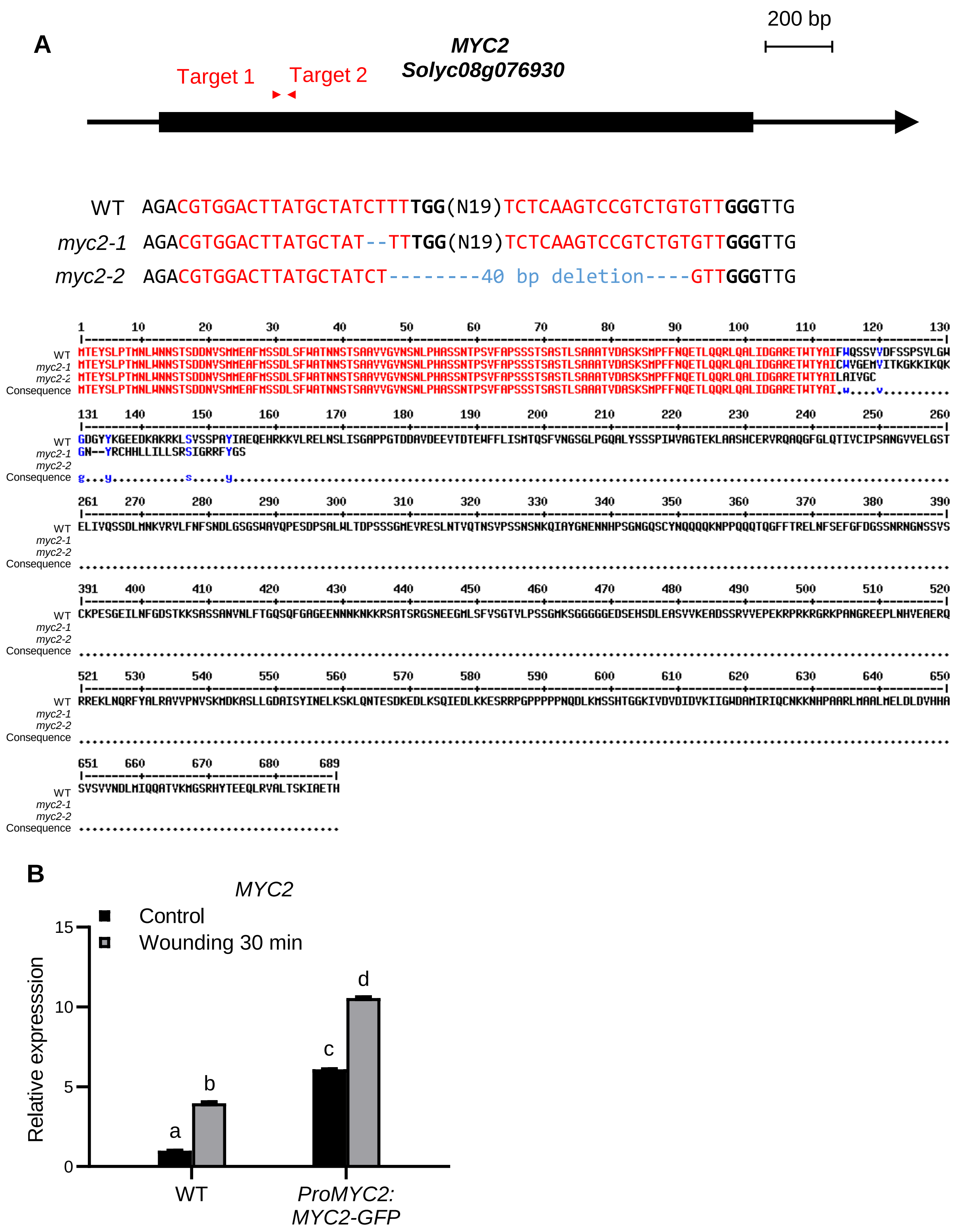


**Supplemental Figure 5. Generation of *myc2* mutants and *ProMYC2:MYC2-GFP* transgenic plants. Related to Figure 5.**

**(A)** Generation of *myc2* mutants. Upper panel: schematic representation of *MYC2* showing the design of CRISPR/Cas9-mediated gene editing. Middle panel: DNA sequence analysis of *myc2* null alleles generated by gene editing. Bottom panel: predicted amino acid sequences encoded by *myc2* null alleles.

**(B)** RT-qPCR results showing wound-induced *MYC2* expression in WT and *ProMYC2:MYC2-GFP* plants. Data are mean ± SD, $n$ = 3 repeats. The bars with different letters are significantly different from each other ($P < 0.001$; Student's $t$ test).
